# Local variation in brain temperature explains gender-specificity of working memory performance

**DOI:** 10.1101/2024.02.14.580344

**Authors:** Jacek Rogala, Joanna Dreszer, Marcin Sińczuk, Łukasz Miciuk, Ewa Piątkowska-Janko, Piotr Bogorodzki, Tomasz Wolak, Andrzej Wróbel, Marek Konarzewski

## Abstract

Exploring gender differences in cognitive abilities offers vital insights into human brain functioning. Our study utilized advanced techniques like magnetic resonance thermometry, standard working memory n-back tasks, and functional MRI to investigate if gender-based variations in brain temperature correlate with distinct neuronal responses and working memory capabilities. Interestingly, our findings revealed no gender disparity in working memory performance. However, we observed a significant decrease in average brain temperature in males during working memory tasks, a phenomenon not seen in females. Although changes in female brain temperature were not statistically significant, we found an inverse relationship between the absolute temperature change (ATC) and cognitive performance, alongside a correlation with blood oxygen level dependent (BOLD) neuronal responses. This suggests that in females, ATC is a crucial determinant for the link between cognitive performance and BOLD responses, a linkage not evident in males. Our results also suggest that females compensate for their brain’s heightened temperature sensitivity by activating additional neuronal networks to support working memory. This study not only underscores the complexity of gender differences in cognitive processing but also opens new avenues for understanding how temperature fluctuations influence brain functionality.

**Significance:** Sex/gender differences in cognition are of high scientific and social interest. Yet, those differences (if any) remain elusive. Here we used magnetic resonance thermometry and functional MRI to examine, whether gender differences in working memory performance (WMP) are determined by subtle, yet detectable between-sex differences in local brain temperature fluctuations mediated by blood oxygen level-dependent (BOLD) neuronal responses. We found that WMP did not differ between genders. Yet, a female’s WMP was more sensitive to brain temperature variation compared to males. Furthermore, the negative impact of temperature on female cognitive functions was compensated by higher BOLD activity in other task-specific brain areas. This compensation may account for equivocal results of studies on the between-sex differences in cognitive performance.

## Introduction

The cognitive abilities of men and women have been under scientific scrutiny for over a century, with a key area of interest being working memory. Playing a crucial role in brain information transfer, working memory affects other cognitive domains, implying that a gender difference in working memory could have broader impacts on cognitive functioning. Yet, behavioral investigations into the influences of gender have yielded inconsistent results. Some meta-analyses suggest a female advantage in verbal working memory (Asperholm et al., 2019; Hirnstein et al., 2022), while others demonstrate no performance differences (Voyer et al., 2017), a female disadvantage (Zilles et al., 2016), or show mixed results pointing to an advantage in males or females depending on the working memory task (Voyer et al. 2021).

Sexual dimorphism in working memory performance appears to be functionally linked to underlying neurophysiology. Previous research has identified sex-based differences in cerebral blood flow (Aanerud et al., 2017, Esposito et al., 1996, Daniel et al., 1989, Rodriguez et al., 1988) and synaptic strength (Aanerud et al., 2017). Remarkably, even when no performance differences between genders are reported, distinct brain activation patterns persist in males and females (Alarcón et al. 2014). A plausible explanation is the variation of brain temperature resulting from obligatory heat cogeneration by metabolic processes underlying neuronal activation. Given that temperature impacts all physical and biochemical processes at the rate of the Van’t Hoff coefficient (McCullough et al., 1999), even small variations, less than one degree Celsius, might adversely affect neuronal activity (Kim et al. 2022). Importantly, the observed daily variations in human brain temperature often exceeded four degrees Celsius (Rzechorzek et al. 2022), and, therefore, may significantly affect neuronal cognitive processes.

Interestingly, the main factors driving changes in brain temperature are neuronal activity and the cooling effect of incoming blood (Kiyatkin et al., 2002). Therefore, it’s plausible, based on existing literature that changes in brain temperature could differ across genders and differentially affect male and female cognitive performance and overall brain activity. However, research into this specific area is still in its infancy, largely because investigations involving human brain temperature typically require invasive techniques. These procedures present ethical considerations and potential risks to patients, such as damage to nervous tissues, making them unsuitable for studies exploring brain activity or cognitive performance.

In theory, one could estimate brain temperature’s effects on cognitive performance using core body temperature as a surrogate. However, the correlation between core body and brain temperature in humans is modest (Thrippleton et al., 2014). Therefore, attempting to infer brain temperatures from core body temperatures could lead to considerable inaccuracies. A more appropriate approach for healthy individuals is to employ non-invasive techniques accurate in examining three-dimensional objects with high water content, such as biological tissues. Magnetic resonance spectroscopic thermometry (MRS-t) is one such technique, which uses magnetic resonance imaging (MRI) scanners to measure temperature-dependent nuclear magnetic resonance (NMR) parameters. The same MRI scanners used for temperature estimation can be used for functional MRI (fMRI) brain imaging using the blood oxygen level dependent (BOLD) technique. The BOLD method evaluates regional differences in cerebral blood flow to ascertain regional brain activity, thereby providing essential information for studying human brain functions. Simultaneous fMRI and electrophysiological recordings have validated that BOLD signaling reflects aspects of neural responses elicited by a stimulus in relevant brain areas (Logothetis & Wandell, 2004). MRS-t and fMRI can be smoothly integrated within a single fMRI session to decode the interplay between gender, brain temperature variations, and brain activity. Continuous advancements over the past three decades have increased the accuracy and reliability of MRS-t, as confirmed by numerous studies (Sung et al., 2022; Winter et al., 2016; Lutz et al., 2020; Zhang et al., 2020).

In our study, we investigated whether gender-specific variations in brain temperatures induced by a working memory task could explain the different neuronal responses and performances observed in men and women. More specifically, we predicted that task-induced variations in brain temperature may drive gender-specific associations between cognitive performance and rates of metabolic processes in activated brain regions. To explore these associations, we combined n-back tasks (a widely used and thoroughly studied experimental cognitive paradigm, Finc et al., 2017; Gevins et al., 1993) with MRS-t to estimate variations in brain temperature, and used fMRI to record activity in 16 brain regions of interest (ROIs) involved with the task during a single fMRI session.

## Results

### Brain temperature and Absolute Temperature Difference (ATC)

Initial (TB_start_) and final (TB_end_) brain temperatures, measured in the right parietal lobe, obtained from the fMRI session averaged 37.32 °C (n = 64 (63); SD = 0.50; min/max: 36.12, 38.62) and 37.07 °C (SD = 0.57, min/max: 35.78, 38.25), respectively. TB_end_ was significantly lower than the TB_start_ (F(1,61) = 18.70, p = < 0.001, η_p_^2^ = 0.244). The repeated measures ANOVA for the difference between TB_start_ and TB_end_ yielded significant brain temperature * sex interaction (F(1,61) = 4.37, p = 0.041, η_p_^2^ = 0.067). Subsequent post hoc tests showed a significant decrease in brain temperatures in men (n = 31; TB_start_: M = 37.37, SD = 0.49, TB_end_: M = 36.98, SD = 0.55), but not in women (n = 32; TB_start_: M = 37.28, SD = 0.53, TB_end_: M = 37.14, SD = 0.59, Figure 1).

**Figure 1.**
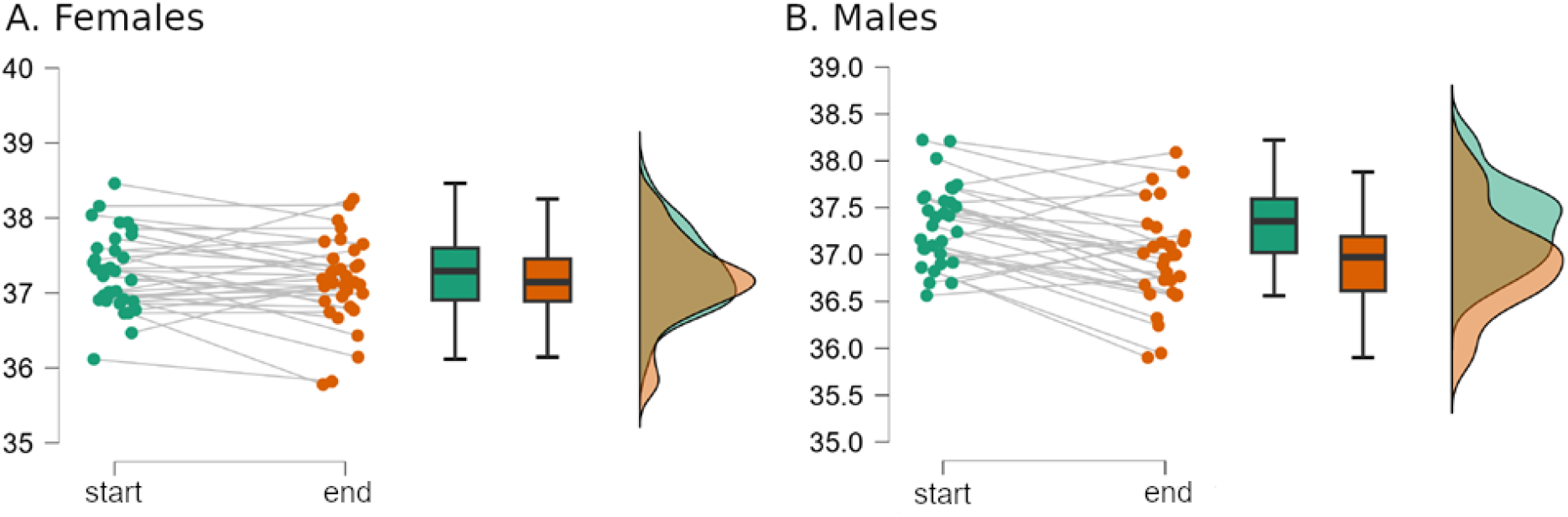
Brain temperature in females and males at the beginning (start) and at the end (end) of an MRI session. Repeated measures ANOVA and subsequent post hoc tests showed significant decreases in brain temperatures in men (n = 31; TB_start_: M = 37.37, SD = 0.49, TB_end_: M = 36.98, SD = 0.55; p < 0.001), but not in women (n = 32; TB_start_: M = 37.28, SD = 0.53, TB_end_: M = 37.14, SD = 0.59; p > 0.1).

Both increases and decreases in temperature can affect neuronal activity. To take into consideration this bi-directionality effect of temperatures, we used the absolute difference between the initial and final brain temperature changes (ATC). This approach, detailed in the materials and methods section, will be consistently applied throughout this paper.

The mean of the ATC of the right inferior posterior lobule was significantly higher in males (0.54 °C) than in females (0.38 °C), t = -2.340, p = 0.020, Cohen’s d = -0.604 (Figure 2).

**Figure 2.**
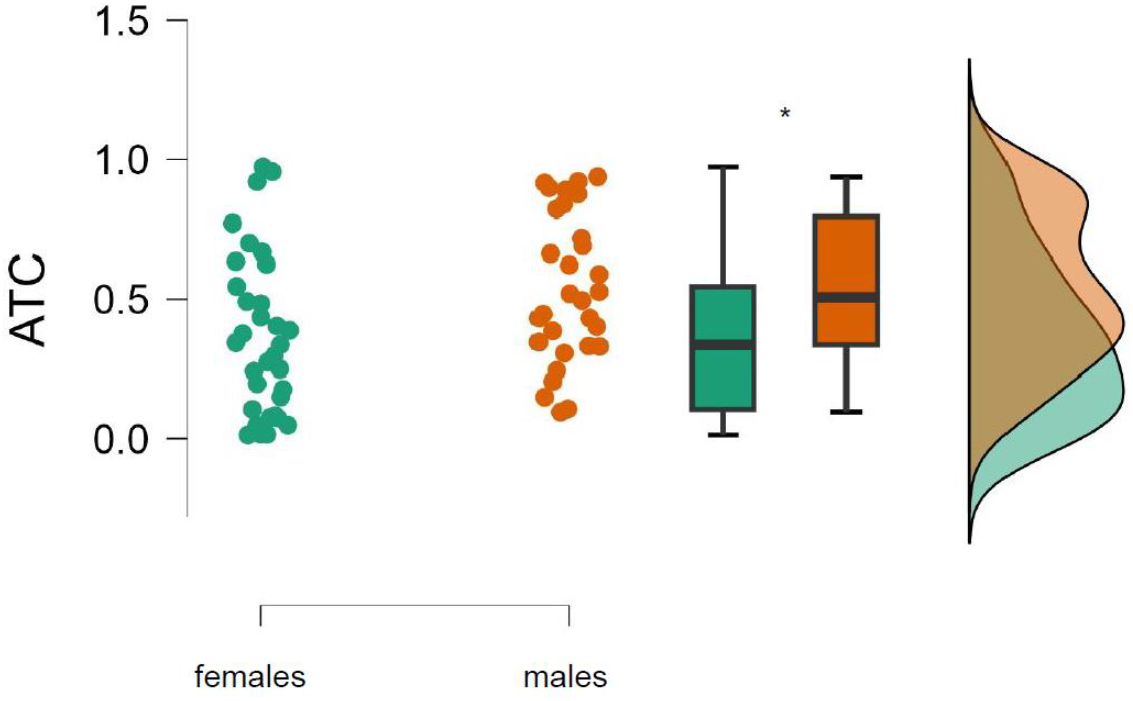
Absolute brain temperature change (ATC) in the right inferior posterior lobule among males (n = 31) and females (n = 32) during a 1-hour MRS/fMRI session. *Significant difference (p = 0.02).

### Distinct effects of brain temperature on gender performance on n-back tasks

To explore the potential interactions between gender and ACT on memory performance, we conducted a series of hierarchical regression analyses. The interaction (model 1, PROCESS macro, Hayes, 2022; see the materials and methods for details) between gender and ATC on predicting working memory performance (measured by d prime index, for details please refer to the behavioral performance evaluation in the materials and methods section) was significant only in the most challenging 2-back testing conditions (F(3,59) = 4.14, p = 0.01, R^2^ = 0.17, B = 0.743, SE = 0.348, 95%, CI = [0.05, 1.43], t = 2.15, p = 0.04). Interestingly, the association of ATC and d prime was negative in women (simple effect test, slope = -0.7936, SE = 0.2322, 95%, CI = [-1.25, -0.32], t = -3.45, p = 0.0011, Figure 3A), while in men the association was insignificant (simple effect test, slope = -0.054, SE = 0.25, 95%, CI = [-0.56, 0.46], t = -0.2122, p = 0.843, Figure 3B).

**Figure 3.**
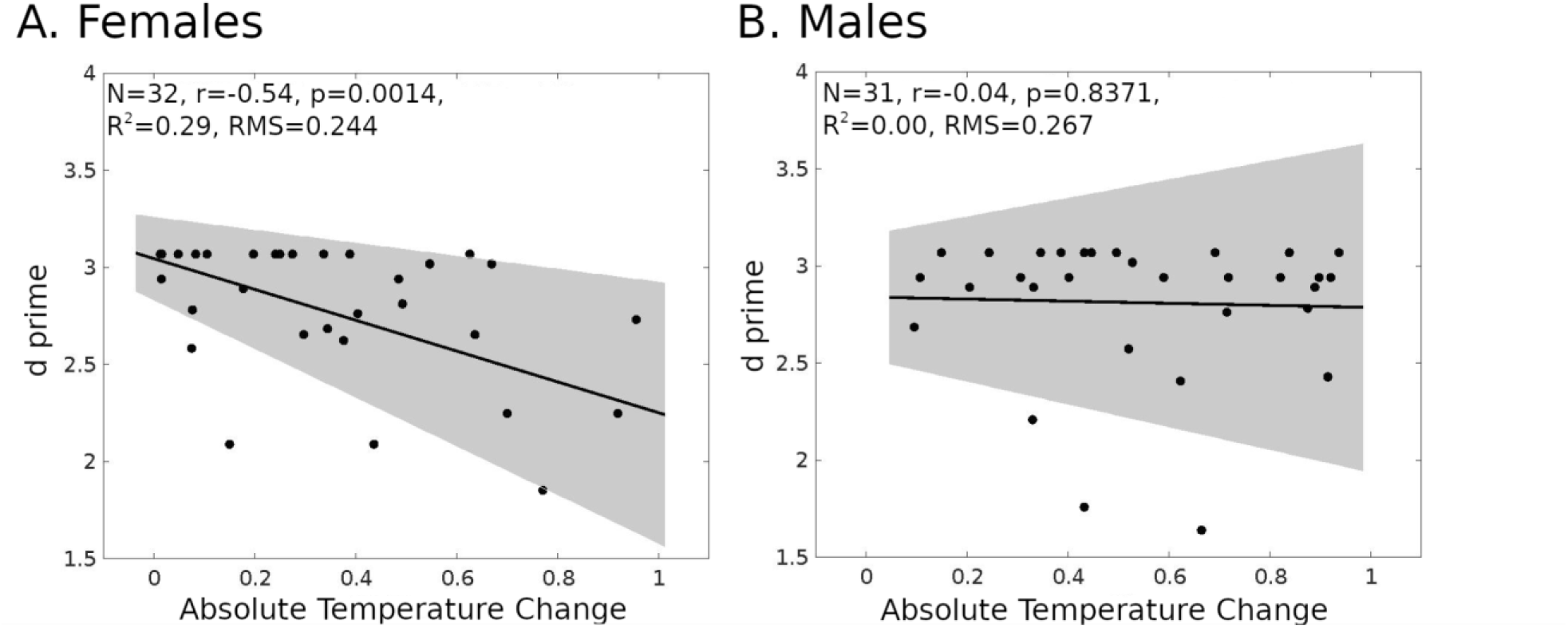
Linear regression models for d prime and absolute temperature change (ATC). A. Female (n = 32). B. Male (n = 31). The shaded gray region is the 95% confidence level of the regression line.

In the less demanding 1-back task, variations had no effect on ATC and working memory performance in males, females, and the whole group.

Most surprisingly, however, despite gender-dependent differences in ATC and ATC-dependent task performance, the behavioral outcomes of both genders were the same (Wilcoxon test Z = - 0.34, p = 0.73). The most plausible explanation for this could be gender- and ATC-dependent differences in brain activity. To test this hypothesis, we started with analyses of brain activity in task-related cerebral ROIs.

### Neural correlates of ATC in task-related ROIs

In women, the observed negative regressions between ATC and performance in most cognitively demanding 2-back task variants suggested a potentially similar association between ATC and BOLD activity. Therefore, we analyzed the regressions between ATC and BOLD in 1-back and 2- back task variants for visual-verbal stimuli in the ROIs (Figure 4). These analyses (FDR corrected) yielded significant results only in the more demanding 2-back task variants. We found a negative relationship between ATC and BOLD in the left ventrolateral prefrontal cortex (left VLPC, adjusted R^2^ = 0.4696, p = 0.0004), the left premotor cortex (left PMC, adjusted R^2^ = 0.2295, p = 0.0289), the right premotor cortex (right PMC, adjusted R^2^ = 0.2227, p = 0.0289), and the left inferior parietal lobule (left IPL, adjusted R^2^ = 0.2856, p = 0.0231; Figure 4). Identical analyses conducted for males did not yield significant results.

**Figure 4.**
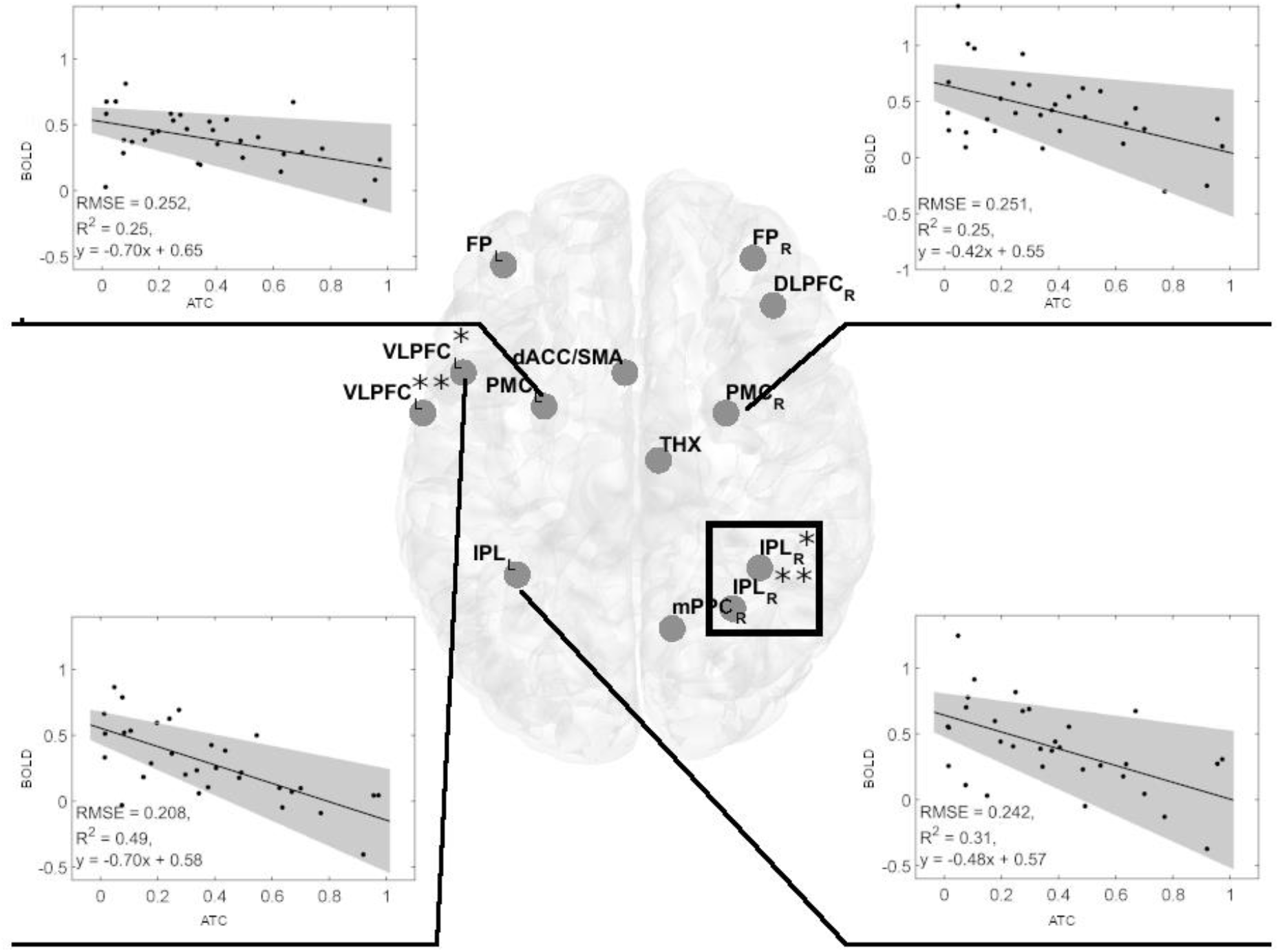
Significant negative linear regression models of BOLD activity in cerebral regions of interest (ROIs) for the visual-verbal 2-back tasks and absolute brain temperature changes in females. Linear regression analyses showed four significant models in the 2-back vs. 1-back comparison (indicated by black lines). Inserts show the significant linear regression models for BOLD changes and ACT. All models were significant at p < 0.05, FDR corrected. The shaded gray region is the 95% confidence level of the regression line. At VLPC: *denotes frontal inferior operculum, and **denotes Rolandic operculum. At IPL: *denotes inferior parietal gyrus, and **denotes anterior cingulum. The black rectangle denotes the spectroscopy voxel of interest where brain temperature was estimated.

Abbreviations: FP_L,R 0_ - frontal poles, VLPFC_L_ - left ventrolateral prefrontal cortex (frontal inferior operculum and Rolandic operculum), dACC_L_ - left dorsal anterior cingulate cortex, DLPFC_R_ - right dorsolateral prefrontal cortex, PMC_L,R_ - bilateral premotor cortex, THX_R_ - right thalamus, IPL_L_ - left inferior parietal lobule, IPL_R_ - two sites in the right inferior parietal lobe, mPPC_R_ - the medial posterior parietal cortex.

An inverse relationship between d prime and ATC (Figure 3), along with an inverse relationship between BOLD and ATC (Figure 4), suggests that both relationships lead to a positive relationship between d prime and BOLD, driven by ATC. Indeed, in women, we observed a positive BOLD-d prime relationship in following ROIs: left VLPFC (adjusted R^2^ = 0.285, p = 0.012, FDR corrected), left PMC (adjusted R^2^ = 0.273, p = 0.012, FDR corrected), and left IPL (adjusted R^2^ = 0.228, p = 0.0203, FDR corrected, Figure 5). The fourth ROI was located in the right medial and lateral cerebellum (adjusted R^2^ = 0.176, p = 0.0432, FDR corrected, Figure 5). Conversely, correlation analyses performed separately for men found none of the correlations to be significant.

**Figure 5.**
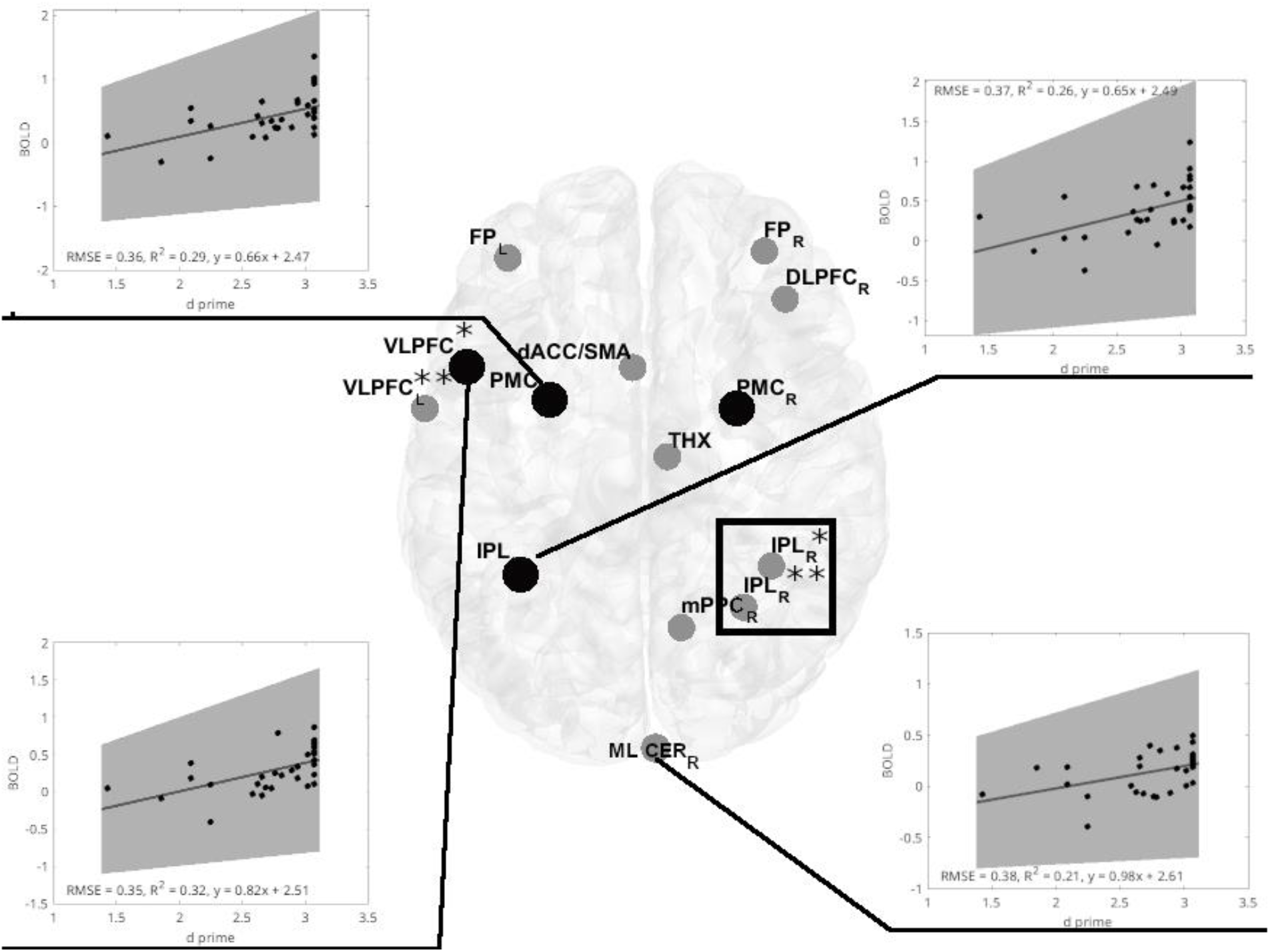
Significant positive linear regression models of d prime vs. BOLD activity in cerebral regions of interest (ROIs) for the visual-verbal 2-back task in females. Linear regression analyses show four significant models in the 2-back vs. 1-back comparisons (indicated by black lines). Out of four significant models, three are located in the same ROIs, identifying an inverse BOLD-ATC relationship (marked with black-filled circles). Inserts show significant linear regression models for BOLD and d prime. All models were significant at p < 0.05, FDR corrected. The shaded gray region is the 95% confidence level of the regression line. At VLPC: *denotes frontal inferior operculum, and **denotes Rolandic operculum. At IPL: *denotes inferior parietal gyrus, and **denotes anterior cingulum. The black rectangle denotes the spectroscopy voxel of interest where brain temperature was estimated.

Abbreviations: FP_L,R 0_ - frontal poles, VLPFC_L_ - left ventrolateral prefrontal cortex (frontal inferior operculum and Rolandic operculum), dACC_L_ - left dorsal anterior cingulate cortex, DLPFC_R_ - right dorsolateral prefrontal cortex, PMC_L,R_ - bilateral premotor cortex, THX_R_ - right thalamus, IPL_L_ - left inferior parietal lobule, IPL_R_ - two sites in the right inferior parietal lobe, mPPC_R_ - the medial posterior parietal cortex, ML CER_R_ - R-Medial and lateral cerebellum.

Further comparisons of BOLD activity between male and female ROIs demonstrated significant differences regardless of the task variant. *Post hoc* tests revealed significantly higher activations for males in five ROIs for the 2-back task variant (Greenhouse-Geisser corrected: F(_7.4,451.7_) = 2.319, p = 0.022, η_p_ ^2^ = 0.037), including three showing significant negative correlations with ATC (Figure 6) and two in the 1-back variant.

**Figure 6.**
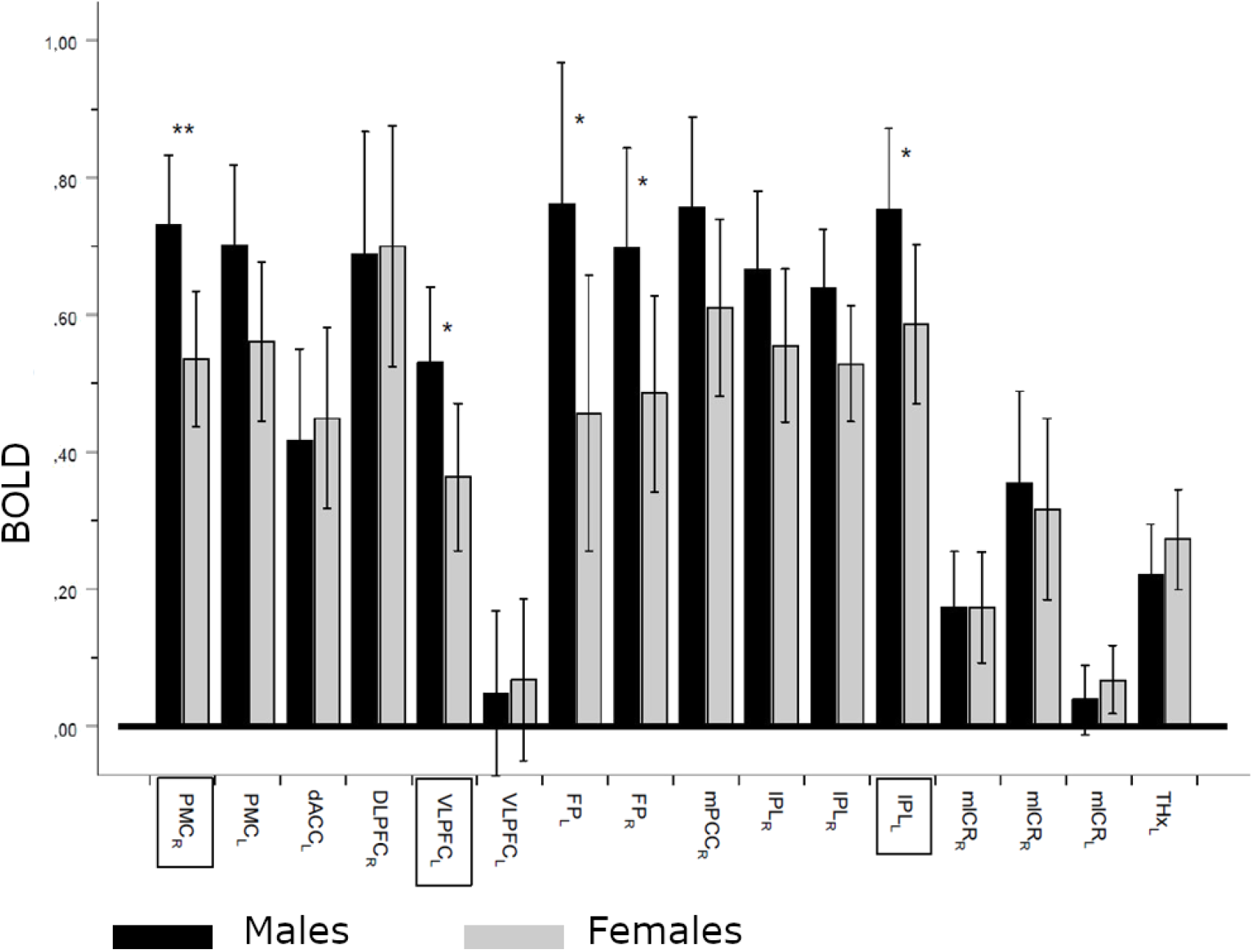
BOLD differences between males and females in regions of interest (ROIs). ROI x sex effect, regardless of the contrast: Greenhouse-Geisser corrected, F(7.405,451,726) = 2.319, p = 0.022, ηp2 = 0.037; *denotes significance level p < 0.05 and **denotes significance level p < 0.01. Dark bars denote means ± CI (95%) of BOLD change for males, and lighter bars denote females. ROI names in frames denote regions with significant negative correlations with absolute temperature differences. PMC_rl_ – bilateral premotor cortex, dACC_l_ – left dorsal anterior cingulate cortex, DLPFC_r_ – right dorsolateral prefrontal cortex, VLPFC_l_ – left ventrolateral prefrontal cortex (frontal inferior operculum and rolandic operculum), FPl_lr_ – left and right frontal poles, mPPC_r_ - right medial posterior parietal cortex, IPL_lr_ – left and right inferior parietal lobule (inferior parietal and anterior cingulum, mlCR_rl_ - medial and lateral cerebellum, THX_r_ – right thalamus.

The above results suggest that women are particularly sensitive to changes in brain temperature, due to decreased BOLD activity in the examined ROIs. However, females maintain comparable working memory performance compared to men despite their pronounced sensitivity to even minor changes in brain temperature. We hypothesize that women compensate for temperature-dependent reductions in BOLD activity in task-related ROIs by activating neuronal activity in other regions supporting working memory. To verify this hypothesis, we compared the whole brain activity of men and women during 2-back and 1-back task variants.

### Whole brain BOLD activity differences between men and women performing n-back tasks

While the comparison of men vs. women (one-sided t-test) whole brain BOLD activity for the 1-vs. 0-back contrast yielded no significant results, the more demanding 2-vs. 1-back contrast showed significant differences in women compared to men. Women showed higher activity in the right inferior parietal lobule (rIPL), right postcentral gyrus (rPoG), right anterior cingulate gyrus (rAnCg), right transverse temporal gyrus (rTTG), and right medial frontal gyrus (rMedFG), Figure 7.

**Figure 7.**
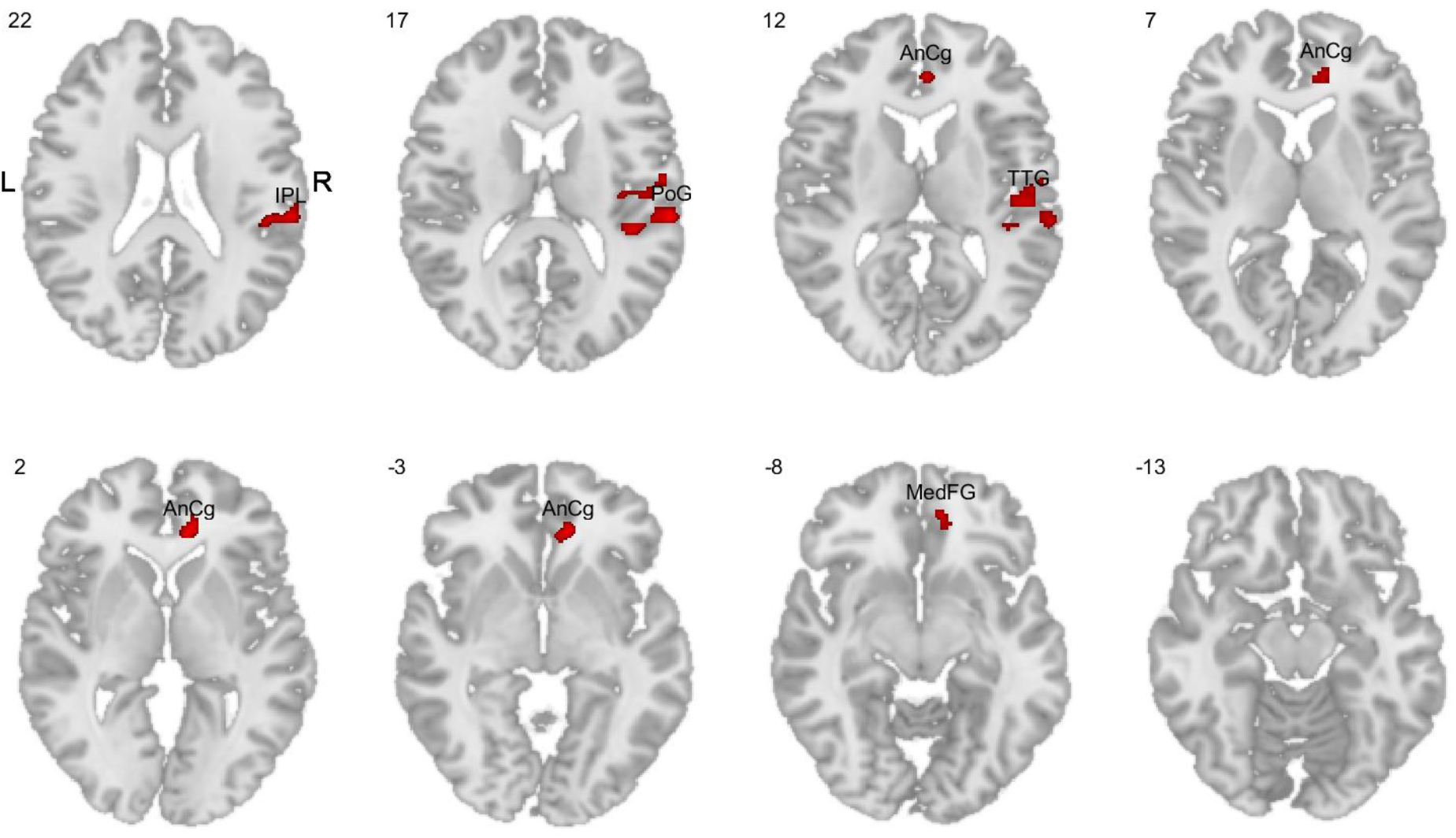
Comparisons of whole brain BOLD activity in the 2-back task variant (1-back vs. 2-back contrast). The red color indicates higher activity in women compared to men. IPL - inferior parietal lobule, PoG - postcentral gyrus, AnCg - anterior cingulate gyrus, TTG - transverse temporal gyrus, MedFG - medial frontal gyrus.

Subsequent analyses did not show significant correlations between BOLD activity and ATC in the above regions. The reversed comparison between men and women did not yield significant results in either task variant or contrast, indicating a lack of male-specific activations.

## Discussion

Our results revealed that commonly observed significant reductions in mean brain temperatures (Sung et al. 2023, Thrippleton et al. 2014, Yablonskiy et al. 2000) were predominantly attributable to male participants, while female brain temperatures remained stable. Despite statistically insignificant changes in female brain temperatures, its variation, evaluated as the ATC, was inversely associated with cognitive performance, as was BOLD, revealing a high sensitivity of the female brain to temperature variations. This strongly suggests that ATC is a primary factor driving the positive association between cognitive performance and BOLD observed in females. Furthermore, our results indicate that the high sensitivity of the female brain to variations in brain temperature is compensated by the activation of additional neuronal regions supporting working memory.

Our study focused on brain areas identified in a meta-analysis as relevant for the execution of n-back tasks (Owen et al. 2005). Although our findings corroborated the involvement of these regions in tasks for the entire participant group, the results were predominantly driven by females. In contrast, no correlations between BOLD activity in the selected ROIs and task performance were observed in males. Therefore, changes in brain temperature adversely affecting a woman’s neural activity, as shown in our study, could affect their behavioral performance, while changes in brain temperature or BOLD activity in men may have no performance consequences.

The lack of correlation between male BOLD activity in task-related regions and ATC, as well as between ATC and task performance, may be due to a man’s generally higher BOLD activity, a finding supported by our results (Figure 6) and previous studies (Dylan et al. 2021, Domes et al. 2010, Sacher et al. 2013). This higher BOLD activity in men could buffer the effects of temperature variations on brain function and performance. In contrast, female brain regions sensitive to ATC showed lower activity, suggesting that lower BOLD levels could lead to reduced neuronal activity due to temperature changes. This may also explain the absence of correlations between BOLD activity and ATC in the less demanding 1-back tasks, which typically show an increase in BOLD activity – a pattern observed in various tasks (Linden et al. 2003, Beauchamp et al. 2001, Callicott et al. 1999). Thus, the observed lack of correlation between ATC and BOLD activity, both in males and in less demanding variants of the tasks in females, may be due to higher BOLD activity mitigating the effect of ATC on neuronal activity.

However, an important finding of our study is that despite a woman’s apparent sensitivity to ATC, their working memory performance remained at the same level as men. One explanation might be that female-specific BOLD activations, like those we detected, support task-specific areas impacted by temperature changes. For instance, our study uniquely identified inferior parietal lobule activation in women, a region implicated in attention and inhibitory processes integral to working memory (Wang et al., 2016, Awh et al., 2006, and others). Another area exclusive to female activation, the medial frontal cortex, is pivotal for memory retention during delay periods and influences working memory capacity via interactions with the dorsolateral prefrontal cortex (Keller et al., 2015). Notably, the most pronounced female-specific BOLD activity was observed in the anterior cingulate cortex (ACC). The ACC is characterized by dual functions, the rostral ACC is involved in overseeing executive cognitive control, while the ventral ACC interacts with emotional processing centers (Bush et al. 2000). During the 2-back tasks, female-specific activations in the ACC also appeared to affect its ventral regions, thereby inducing additional cognitive and emotional processes accompanying task performance. These additional emotional processes are sometimes referred to as characteristic problem-solving strategies employed by women (see Hill et al. 2014 for a review), whereas in light of our results, they are likely a side effect of compensatory activations of the ACC during the n-back tasks.

MRS-t is relatively new and is validated against invasive methods. Studies using invasive human brain temperature measurements performed on patients with acute brain injury indicate that the brain temperature ranges between 32.6 °C and 45.0 °C with average daily and hourly variations within one degree (Rossi et al., 2001, Lu et al., 2021, Rzechorzek et al., 2022). However, individual variations during 72-hour monitoring reach as high as 4 degrees depending on the patient’s behavioral state (Rzechorzek et al., 2022). Our measurements ranged from 35.77–38.61 °C with an amplitude of 0.45 degrees, which falls in the middle of invasive results. They also fall within the range of other MRS-t measurements (Zhang et al., 2020: 35.4–37.4 °C; Rzechorzek et al., 2022: 36.1–40.9 °C; Sung et al., 2022: 35.5–40.1 °C). Thus, it can be concluded that temperature measurements in our study are within the range of amplitudes for both invasive and non-invasive readings obtained in other investigations and can be considered a reliable estimate of brain temperature.

In conclusion, our results revealed that a woman’s performance in cognitive tasks is highly sensitive to variations in brain temperature. Female brain temperature-related decreases in performance were compensated by the activation of additional brain regions, not engaged in men. Overall, our study revealed subtle yet important gender-specific differences in neurophysiological mechanisms underlying compensatory mechanisms, yielding the lack of appreciable differences between men and women subjected to cognitive tasks.

## Materials and Methods

### Participants

Sixty-three right-handed healthy adults (32 females) took part in this study with a mean age of 28 years (SD = 5.5 years, range = 21–42 years). Participants were paid for their time. They were homogeneous in terms of their socioeconomic status and education level, and the majority were students. Following the ethical principles for medical research involving human subjects, ethical approval was granted by the local ethics committee, and all participants provided informed consent after a thorough explanation of the study’s procedures and risks. Participants were included in this experiment after verifying the absence of contraindications to MRI examinations (e.g., claustrophobia, cochlear implants, metal fragments in the participants’ eye or head, pacemakers) as well as other conditions posing a potential risk for the participants.

### Acquisition of MRI data

MRI data acquisition was carried out using a 3-T GE Discovery 750w scanner (CNS Lab IBBE PAS) fitted with a standard receiver, 8-channel head coil. Room temperature (20.5 °C, standard deviation [SD] ± 0.5 °C) and lighting were constant. After standard shimming, a high-resolution structural MRI sequence (TR/TE = 6.936 ms/2.968 ms, matrix 256 × 256, voxel 1.0547 × 1.0547 × 1.2 mm3) was acquired for improved voxel localization. A single-voxel 1H-MRS spectra (PRESS) was acquired from a voxel of interest (VOI) (20 × 20 × 20 mm3) positioned in the right inferior parietal lobule (rIPL). This large VOI ensures a reasonable signal-to-noise ratio to gain sufficient 1H-MRS signal for further data processing. It is also one of the regions of interest (ROIs) identified by Owen and colleagues (2005) in their meta-analysis devoted to n-back working memory tasks. We used single voxel spectroscopy (SVS) for its high signal-to-noise ratio during a relatively short scan time, and the compact imaging area allowed shimming with high-quality spectra suitable for peak parameterization allowing higher precision of temperature estimation than spectroscopy imaging methods (Bertoldo et al. 2013). The scan parameters for the PRESS were as follows: TR = 1500 ms, TE = 30 ms, bandwidth 5 kHz, 4096 points, NEX = 8, with 96 averages and a scan time of 3 minutes. For more precise data analysis, water suppression during the scanning sequence was applied. After MRS scanning, six functional scans were collected using a t2*-weighted gradient echo planar imaging sequence (TR = 2500 ms, TE = 25 ms, voxel 3 × 3 × 3 mm3). During the first two scans, lasting 5 minutes and 50 seconds, participants performed an Anticipation Attention Task, whereas the last four scans each lasted 4 minutes and 25 seconds while n-back tasks (Finc et al., 2017) were carried out. After functional scans, a second pair of PRESS scans were conducted using the same parameters.

### In vivo temperature measurements

All measurements were acquired between 8:00 am and 4:00 pm. The body temperature of the participants was recorded three times before the examination using an infrared ear thermometer (Braun ThermoScan® 3) in the right ear. The participants were fitted comfortably inside the scanner and equipped with earplugs, their head was positioned centrally within the coil, and supported by padding and foam to ensure minimal movement in any plane. Subsequently, the acquisition of the data was performed (see Acquisition of MRS data section for detailed description). On completion of the scan, three additional temperature measurements in the right ear were obtained.

The total time scheduled was approximately 60 minutes for each participant: fifteen minutes for participant preparation and 45 minutes for MRS acquisition. A total of 128 MRS measurements were performed (64 participants, each with 2 PRESS scans where each was composed of water-suppressed and unsuppressed data).

### MRS data preprocessing and analysis

The preprocessing and temperature estimations were performed using house-developed automated software based on FID-A (Jamie Near, McGill University, 2014), an open-source software package written under a Matlab™ platform (MathWorks, Natick, MA, USA). After the preprocessing of the MRS spectra (Near et al., 2021), covering the following steps: Fourier transformation, RF coil combination, signal averaging, and phase correction localization of individual metabolites and water peaks was done with Lorentzian fitting. Brain temperature was estimated using the following equation:

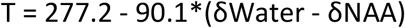

where:

T - is the temperature (°C),

δWater - is the chemical shift of the H20 peak, and

δNAA - is the chemical shift of the N-acetylaspartate (NAA) peak

The constant coefficients in the equation were measured with a temperature-controlled phantom (Sińczuk et al., 2023) containing: N-acetylaspartate, choline, and creatine metabolites in the proportions as observed in the healthy human brain. The accuracy of temperature estimations, according to past research is within approximately 0.5 °C (Verius et al., 2019).

Due to a short period of temperature monitoring (two readings 1 hour apart), the regular cosinor analysis required to identify circadian temperature rhythm and amplitude had to be adjusted (Lu et al., 2021). We used absolute temperature differences as the proxy for amplitude.

### Absolute Temperature Change (ATC)

Because each cellular and molecular process is likely to have a different temperature dependence or Q10, predicting neuronal activity with temperature changes is not trivial. Available reports on temperature-dependent neuronal activity show a plethora of possible responses, for instance, Masino et al. found that increasing the temperature markedly inhibits excitatory synaptic transmissions, while other investigations showed that lower temperatures caused slower nerve conduction velocities and increased nerve potential amplitudes (Denys 1991). Yet, another study indicates that field excitatory potentials were linearly related to brain temperature (Moser et al. 1993), while Boulant et al. (1974) found both negative and positive coefficients for temperatures and firing rates. Thus, to account for differentiated neuronal responses during temperature changes, we adopted ATC as the absolute value of the difference between initial (TBstart) and final (TBend) brain temperatures. Similar measures - brain temperature amplitude – are commonly used in longitudinal brain temperature monitoring in clinical and research practice (Rzechorzek et al., 2022; Kiyatkin et al., 2002).

### fMRI (Blood Oxygen Level Dependent) data preprocessing and analysis

The data preprocessing pipeline was done using the SPM12 toolbox (Wellcome Department of Imaging Neuroscience, Institute of Neurology, London, UK) running on Matlab (R2016a) (Mathworks, Natick, MA, USA). Functional images were corrected for acquisition time and spatially realigned to the mean image to correct for interscan head motion. The structural T1-weighted image was co-registered with the mean functional image. Functional, gray matter, white matter, and CSF images were spatially normalized to the MNI template using new unified normalization segmentation. One participant and five single sessions from other participants were excluded from further analysis due to excessive head motion artifacts (more than 10 in outlier scans), thus, the final sample consisted of 63 subjects. No significant differences in the number of outlier scans were found between 1-back and 2-back tasks (t(63) = 0.38, p = 0.82).

The I-level analysis of the blood oxygen level dependent (BOLD) signals was done under an SPM12 general linear model (GLM) framework. A design matrix consisted of three task-related regressors corresponding to 0-back, 1-back, and 2-back scanning sessions, session-specific effects, and nuisance variables like subject head movement. A linear combination of effect sizes for task regressors (1-vs. 0-back and 2-vs. 1-back) was chosen with contrast vectors for further group-level analysis. On II-level analysis, additional regressors such as initial brain temperature, final brain temperature, change in temperature during the study, and gender were defined. Based on previous publications (Owen et al., 2005), 16 ROIs were selected for which the mean activity values were determined for the two mentioned BOLD contrasts. ROIs identified by Owen and colleagues (2005) devoted to the verbal variant of n-back working memory tasks included the bilateral premotor cortex, left dorsal anterior cingulate cortex, right dorsolateral prefrontal cortex, left ventrolateral prefrontal cortex (frontal inferior operculum and rolandic operculum), left and right frontal poles, right medial posterior parietal cortex, left and rIPL (inferior parietal and anterior cingulum), and three sites in the medial and lateral cerebellum and right thalamus.

### Working memory task

To assess working memory performance, we used classical n-back tasks. Details of the task used in the current study were described in our previous work (Finc et al., 2017). In short, a visual letter-based n-back task (Gevins & Cutillo, 1993) requires participants to continuously monitor, update, and manipulate information held in memory (Owen et al., 2005). In our study, the stimulus consisted of the first five letters of the Latin alphabet (A–E, Kearney-Ramos et al., 2014), presented in pseudorandom order in the 0-, 1-, and 2-back conditions. The presentation of each letter (500 ms) was preceded by a fixation cross (1500 ms) presented in the center of the screen. Each session (RUN) of the task consisted of 10 blocks (30 seconds per block, 12 trials with 25% targets), in which 0-, 1-, and 2-back conditions alternated. Before each block, brief instructions (2 seconds) were displayed to inform the participant about the current task. The task consisted of three RUNs with a total length of 16 minutes; 480 functional scans were performed during each RUN. Depending on the conditions, participants were asked to indicate whether the currently presented letter was the same or different from the letter presented one or two trials earlier. Participants had 2000 ms for each response. The execution and control of the experimental protocol (stimulus delivery and response recording) were performed by Presentation® software (Neurobehavioral Systems, Albany, NY), version 17.2. Visual stimuli were displayed on a monitor and presented to the participant in a mirror. Participants were instructed to respond as quickly and accurately as possible by pressing one of two buttons with the thumb of their right hand (target letter -right button, non-target letter - left button).

### Behavioral performance evaluation

To evaluate a participant’s performance in the n-back tasks, we used the sensitivity index d prime (Macmillan & Creelman et al., 2000). The d prime stands for the difference between Z transforms of the hit rate and false alarms. The hit rate is defined as the proportion of hits when a signal is present to all target stimuli, and false alarms represent the proportion of responses when a signal is absent to all non-target stimuli. Final scores were adjusted according to the following formulas:

1 − 1/(2n) for perfect hits and 1/(2n) for zero false alarms, where n was the number of total hits or false alarms (Macmillan & Creelman et al., 2000).

### Statistical methods

Data were tested for normality using the Anderson-Darling test and checked for the presence of outliers. The values deviating from the mean by more than three standard deviations were removed. Comparison of the groups was conducted using two-tailed, two-sample t-tests or repeated measures ANOVA. In cases of a small number of samples, the Wilcoxon test and Kruskal-Wallis non-parametric ANOVA with Chi-square post-hoc test were used. For correlation analyses, the Pearson correlation and regression analyses with adjustments were applied. Results were corrected for false positives using the False Discovery Rate adjustment (Benjamini & Hochberg 1995) where appropriate unless otherwise stated. Analyses and figures were created using MATLAB 2016a.

To verify whether there is a moderation effect of gender, we applied a series of moderation analyses (Model 1, Hayes, 2022, via SPSS, macro PROCESS v. 3.2), which allowed us to evaluate and compare direct, indirect, and conditional effects. The method recommended by Hayes (2022) does not require the data to fulfill the criteria of normality and it is based on the estimation of bias-corrected, 95% bootstrapped confidence intervals from a series of 10,000 bootstrap samples.

## Declaration of interests

The authors declare no competing interests.

## Funding

National Centre for Research and Development; grant POIR-01.01.01-00-178/15

## Author contributions

Conceptualization: JR, JD

Methodology: JR, MS, TW, EPJ, PB, ŁM, UM

Investigation: MS

Statistical analyses: JD, JR, ŁM

Visualization: TW, JR, JD

Project administration: JR, UM

Supervision: PB, EPJ, AW, MK

Writing – original draft: JR, MS, ŁM, TW

Writing – review & editing: AW, JD, UM, MK

## Competing interests

Authors declare that they have no competing interests.

## Data and materials availability

All data available on request.

## Notes

### Competing Interest Statement

The authors have declared no competing interest.

## References

1. Aanerud, J., Borghammer, P., Rodell, A., Jonsdottir, K. Y., & Gjedde, A. (2017). Sex differences of human cortical blood flow and energy metabolism. Journal of Cerebral Blood Flow & Metabolism, 37(7), 2433–2440.

2. Aronov D, Fee MS (2012) Natural Changes in Brain Temperature Underlie Variations in Song Tempo during a Mating Behavior. PLoS ONE 7(10): e47856. 10.1371/journal.pone.0047856

3. Babiloni C, Noce G, Ferri R, Lizio R, Lopez S, Lorenzo I, Tucci F, Soricelli A, Zurrón M, Díaz F, Nobili F, Arnaldi D, Famà F, Buttinelli C, Giubilei F, Cipollini V, Marizzoni M, Güntekin B, Yıldırım E, Hanoğlu L, … Del Percio C (2022) Resting State Alpha Electroencephalographic Rhythms Are Affected by Sex in Cognitively Unimpaired Seniors and Patients with Alzheimer’s Disease and Amnesic Mild Cognitive Impairment: A Retrospective and Exploratory Study. Cerebral cortex 32(10), 2197–2215. doi:10.1093/cercor/bhab348

4. Badre D, Poldrack RA, Pare-Blagoev EJ, Insler RZ, Wagner AD (2005) Dissociable Controlled Retrieval and Generalized Selection Mechanisms in Ventrolateral Prefrontal Cortex, Neuron 47, 907–918. doi: 10.1016/j.neuron.2005.07.023

5. Beauchamp, M. S., Petit, L., Ellmore, T. M., Ingeholm, J., & Haxby, J. V. (2001). A parametric fMRI study of overt and covert shifts of visuospatial attention. Neuroimage, 14(2), 310–321.

6. Benjamini Y. & Hochberg Y. Controlling the False Discovery Rate: A Practical and Powerful Approach to Multiple Testing. Journal of the Royal Statistical Society B. 57, 289–300 (1995).

7. Bertoldo D, Watcharakorn A, Castillo M (2013) Brain proton magnetic resonance spectroscopy. Introduction and overview. Neuroimag Clin N Am. 23:359–380.

8. Boulant, J. A., & Hardy, J. D. (1974). The effect of spinal and skin temperatures on the firing rate and thermosensitivity of preoptic neurones. The Journal of physiology, 240(3), 639.

9. Bush G, Luu P, Posner MI. Cognitive and emotional influences in anterior cingulate cortex. Trends Cogn Sci. 2000 Jun;4(6):215–222. doi: 10.1016/s1364-6613(00)01483-2. PMID: 10827444.

10. Busto R, Dietrich WD, Globus MY et al (1987) Small differences in intraischemic brain temperature critically determine the extent of ischemic neuronal injury. J. Cereb. Blood Flow Metab.7, 729–738 doi: 10.1038/jcbfm.1987.127

11. Byrne BM (2001) Structural equation modeling with AMOS: Basic concepts, applications, and programming. Lawrence Erlbaum Associates Publishers

12. Callicott, J. H., Mattay, V. S., Bertolino, A., Finn, K., Coppola, R., Frank, J. A., … & Weinberger, D. R. (1999). Physiological characteristics of capacity constraints in working memory as revealed by functional MRI. Cerebral cortex, 9(1), 20–26.

13. Caplan JS, Williams AH, Marder E (2014) Many parameter sets in a multicompartment model oscillator are robust to temperature perturbations. J. Neurosci. 34, 4963–4975.

14. Childs C, Hiltunen Y, Vidyasagar R, Kauppinen RA (2007) Determination of regional brain temperature using proton magnetic resonance spectroscopy to assess brain-body temperature differences in healthy human subjects. Magnetic resonance in medicine, 57(1), 59–66. 10.1002/mrm.21100

15. Csernai M, Borbély S, Kocsis K, Burka D, Fekete Z, Balogh V, Káli S, Emri Z, Barthó P (2019) Dynamics of sleep oscillations is coupled to brain temperature on multiple scales. J. Physiol. 597, 4069–4086. doi: 10.1113/JP277664

16. Daniel, D. G., Mathew, R. J., & Wilson, W. H. (1989). Sex roles and regional cerebral blood flow. Psychiatry research, 27(1), 55–64.

17. DeMaegd ML, Stein W (2020) Temperature-robust activity patterns arise from coordinated axonal Sodium channel properties. PLOS Computational Biology 16, e1008057. doi: 10.1371/journal.pcbi.1008057

18. Denys, E.H. (1991), AAEM minimonograph #14: The influence of temperature in clinical neurophysiology. Muscle Nerve, 14: 795–811. 10.1002/mus.880140902

19. Domes G, Schulze L, Böttger M, Grossmann A, Hauenstein K, Wirtz PH, Heinrichs M, Herpertz SC (2010) The neural correlates of sex differences in emotional reactivity and emotion regulation. Hum. Brain Mapp. 31, 758–769. doi: 10.1002/hbm.20903

20. Dylan S. Spets & Scott D. Slotnick (2021) Are there sex differences in brain activity during long-term memory? A systematic review and fMRI activation likelihood estimation meta-analysis, Cognitive Neuroscience, 12:3–4, 163–173, DOI: 10.1080/17588928.2020.1806810

21. Esposito, G., Van Horn, J. D., Weinberger, D. R., & Berman, K. F. (1996). Gender differences in cerebral blood flow as a function of cognitive state with PET. Journal of Nuclear Medicine, 37(4), 559–564.

22. Finc K, Bonna K, Lewandowska M, Wolak T, Nikadon J, Dreszer J, Duch W, Kühn S (2017) Transition of the functional brain network related to increasing cognitive demands. Hum. Brain Mapp. 38, 3659–3674. doi: 10.1002/hbm.23621

23. Gevins A, Cutillo B (1993) Spatiotemporal dynamics of component processes in human working memory, Electroenceph. Clin. Neurophysiol. 87, 128–143. doi: 10.1016/0013-4694(93)90119-G

24. Goldstein JM, Poldrack R, Anagnoson R, Breiter HC, Makris N (2005) Sex differences in prefrontal cortical brain activity during fMRI of auditory verbal working memory. Neuropsychology 19, 509–519. doi:10.1037/0894-4105.19.4.509

25. Guatteo E, Chung KK, Bowala TK, Bernardi G, Mercuri NB, Lipski J (2005) Temperature sensitivity of dopaminergic neurons of the substantia nigra pars Compacta: involvement of transient receptor potential channels Journal of Neurophysiology 94:3069–3080. 10.1152/jn.00066.2005)

26. Haddad SA, Marder E (2018) Circuit Robustness to Temperature Perturbation Is Altered by Neuromodulators, Neuron 100, 609–623.e3. doi: 10.1016/j.neuron.2018.08.035

27. Hardy JD, Du Bois EF (1938) Basal metabolism, radiation, convection and vaporization at temperatures of 22 to 35°C J Nutr, 15 (1938), pp. 477–497

28. Ibaraki M., Shinohara Y., Nakamura K., Miura S., Kinoshita F., Kinoshita T. (2010). Interindividual variations of cerebral blood flow, oxygen delivery, and metabolism in relation to hemoglobin concentration measured by positron emission tomography in humans. J. Cereb. Blood Flow Metab. 30, 1296–1305. doi: 10.1038/jcbfm.2010.13

29. Jackson DL (2003) Revisiting sample size and number of parameter estimates: Some support for the N:q hypothesis. Structural Equation Modeling 10(1), 128–141.

30. Janssen R (1992) Thermal influences on nervous system function. Neurosci. & Biobehav. Reviews 16, 399–413 DOI: 10.1016/S0149-7634(05)80209-X

31. Kearney-Ramos TE, Fausett JS, Gess JL, Reno A, Peraza J, Kilts CD, James GA (2014) Merging clinical neuropsychology and functional neuroimaging to evaluate the construct validity and neural network engagement of the n-back task. J. Int. Neuropsychol. Soc. 20, 736–750.

32. Kim, T., Kadji, H., Whalen, A. J., Ashourvan, A., Freeman, E., Fried, S. I., … & Schiff, S. J. (2022). Thermal effects on neurons during stimulation of the brain. Journal of neural engineering, 19(5), 056029.

33. Kiyatkin EA (2019) Brain temperature and its role in physiology and pathophysiology: Lessons from 20 years of thermorecording. Temperature. 6, 271–333. doi: 10.1080/23328940.2019.1691896

34. Kiyatkin, E. A., Brown, P. L., & Wise, R. A. (2002). Brain temperature fluctuation: a reflection of functional neural activation. European Journal of Neuroscience, 16(1), 164–168.

35. Kline RB (2015) Principles and Practice of Structural Equation Modeling, 4th Edition, New York: Guilford Publications.

36. Linden, D. E., Bittner, R. A., Muckli, L., Waltz, J. A., Kriegeskorte, N., Goebel, R., … & Munk, M. H. (2003). Cortical capacity constraints for visual working memory: dissociation of fMRI load effects in a fronto-parietal network. Neuroimage, 20(3), 1518–1530.

37. Logothetis, N. K., & Wandell, B. A. (2004). Interpreting the BOLD signal. Annu. Rev. Physiol., 66, 735–769.

38. Long M, Fee M (2008) Using temperature to analyse temporal dynamics in the songbird motor pathway. Nature 456, 189–194. doi: 10.1038/nature07448

39. Lutz NW, Bernard M (2020) Contactless Thermometry by MRI and MRS: Advanced Methods for Thermotherapy and Biomaterials. iScience, 23(10), 101561. 10.1016/j.isci.2020.101561

40. Lu, HY, Huang AP, Kuo LT (2021) Prognostic Value of Circadian Brain Temperature Rhythm in Basal Ganglia Hemorrhage After Surgery. Neurology and therapy, 10(2), 1045–1059. 10.1007/s40120-021-00283-y

41. Macmillan NA, Creelman CD (1990) Response bias: Characteristics of detection theory, threshold theory, and “non-parametric” indexes. Psychol. Bull. 107, 401–413. doi: 10.1037/0033-2909.107.3.401

42. Marder E, Haddad SA, Goeritz ML, Rosenbaum P, Kaspersky T (2015) How can motor systems retain performance over a wide temperature range? Lessons from the crustacean stomatogastric nervous system. J. Comp. Physiol. A, Neuroethology, sensory, neural, and behavioral physiology, 201, 851–856. doi: 10.1007/s00359-014-0975-2

43. Masino SA, Dunwiddie TV (1999) Temperature-Dependent Modulation of Excitatory Transmission in Hippocampal Slices Is Mediated by Extracellular Adenosine. J. Neurosci. 19,1932–1939. doi: 10.1523/JNEUROSCI.19-06-01932.1999

44. McCullough, J. N., Zhang, N., Reich, D. L., Juvonen, T. S., Klein, J. J., Spielvogel, D., … & Griepp, R. B. (1999). Cerebral metabolic suppression during hypothermic circulatory arrest in humans. The Annals of thoracic surgery, 67(6), 1895–1899.

45. Moser E, Mathiesen I, Andersen P (1993) Association between brain temperature and dentate field potentials in exploring and swimming rats. Science (New York, N.Y.), 259(5099), 1324–1326. 10.1126/science.8446900

46. Near J, Harris AD, Juchem C, Kreis R, Marjańska M, Slotboom GÖzJ, Wilson M, Gasparovic C (2021) Preprocessing, analysis and quantification in single-voxel magnetic resonance spectroscopy: experts’ consensus recommendations. NMR Biomed. 34, e4257. doi: 10.1002/nbm.4257. Epub 2020 Feb 21. PMID: 32084297; PMCID: PMC7442593.

47. Owen AM, McMillan KM, Laird AR, Bullmore E (2005) N-back working memory paradigm: a meta-analysis of normative functional neuroimaging studies. Hum. Brain. Mapp. 25, 46–59. doi: 10.1002/hbm.20131

48. Robertson RM, Money TG (2012) Temperature and neuronal circuit function: compensation, tuning and tolerance. Curr. Opin. Neurobiol. 22, 724–734.

49. Rodriguez, G., Warkentin, S., Risberg, J., & Rosadini, G. (1988). Sex differences in regional cerebral blood flow. Journal of Cerebral Blood Flow & Metabolism, 8(6), 783–789.

50. Rosen AD (2001) Nonlinear temperature modulation of sodium channel kinetics in GH3 cells Biochimica Et Biophysica Acta (BBA) - Biomembranes 1511:391–396. 10.1016/S0005-2736(01)00301-7

51. Sacher J, Neumann J, Okon-Singer H, Gotowiec S, Villringer A (2013) Sexual dimorphism in the human brain: evidence from neuroimaging, Magnetic Resonance Imaging 31, 366–375. doi: 10.1016/j.mri.2012.06.007

52. Spets DS, Slotnick SD (2021) Are there sex differences in brain activity during long-term memory? A systematic review and fMRI activation likelihood estimation meta-analysis, Cognitive Neuroscience, 12:3–4, 163–173, DOI: 10.1080/17588928.2020.1806810

53. Simkus CRL, Stricker C (2002) Properties of mEPSCs recorded in layer II neurons of rat barrel cortex The Journal of Physiology 545:509–520. 10.1113/jphysiol.2002.022095

54. Sińczuk, M., Rogala, J., Piątkowska-Janko, E., Bogorodzki, P. (2024). Application of Unsuppressed Water Peaks for MRS Thermometry. In: StrumiŁŁo, P., Klepaczko, A., Strzelecki, M., Bociąga, D. (eds) The Latest Developments and Challenges in Biomedical Engineering. PCBEE 2023. Lecture Notes in Networks and Systems, vol 746. Springer, Cham. 10.1007/978-3-031-38430-1_31

55. Sung D, Risk BB, Wang KJ, Allen JW, Fleischer CC (2022) Resting-State Brain Temperature: Dynamic Fluctuations in Brain Temperature and the Brain-Body Temperature Gradient. Journal of magnetic resonance imaging: JMRI, 10.1002/jmri.28376. Advance online publication. 10.1002/jmri.28376

56. Tang LS, Goeritz ML, Caplan JS, Taylor AL, Fisek M, Marder E (2010) Precise temperature compensation of phase in a rhythmic motor pattern. PLOS Biology 8, e1000469. DOI: 10.1371/journal.pbio.1000469, PMID: 20824168.

57. Thrippleton MJ, Parikh J, Harris BA, Hammer SJ, Semple SIK, Andrews PJD, Wardlaw JM, Marshall I (2014) Reliability of MRSI brain temperature mapping at 1.5 and 3 T. NMR Biomed. 27, 183–190. doi: 10.1002/nbm.3050

58. Tryba AK, Ramirez JM (2004) Hyperthermia modulates respiratory pacemaker bursting properties Journal of Neurophysiology 92:2844–2852. 10.1152/jn.00752.2003

59. Voyer, Daniel, et al. “Sex differences in verbal working memory: A systematic review and meta-analysis.” Psychological bulletin 147.4 (2021): 352.

60. Voyer, D., Voyer, S. D., & Saint-Aubin, J. (2017). Sex differences in visual-spatial working memory: A meta-analysis. Psychonomic bulletin & review, 24, 307–33

61. Watson PL, Weiner JL, Carlen PL (1997) Effects of variations in hippocampal slice preparation protocol on the electrophysiological stability, epileptogenicity and graded hypoxia responses of CA1 neurons. Brain Res., 775, 134–143. doi: 10.1016/S0006-8993(97)00893-7.

62. Werner J (1981) Control aspects of human temperature regulation Automatica, 17, pp. 351–362

63. Winter L, Oberacker E, Paul K, Ji Y, Oezerdem Ghadjar P, Thieme A, Budach V, Wust P, Niendorf T (2016) Magnetic resonance thermometry: Methodology, pitfalls and practical solutions, International Journal of Hyperthermia, 32:1, 63–75, DOI: 10.3109/02656736.2015.1108462

64. Wimber M, Bäuml K-H, Bergström Z, Markopoulos G, Heinze H-J, Richardson-Klavehn (2008) A Neural Markers of Inhibition in Human Memory Retrieval. J Neuroscience 28, 13419–13427. doi: 10.1523/JNEUROSCI.1916-08.2008

65. Wolf RC, Vasic N, Walter H (2006) Differential activation of ventrolateral prefrontal cortex during working memory retrieval, Neuropsychologia 44, 2558–2563. doi: 10.1016/j.neuropsychologia.2006.05.015

66. Vasic N, Lohr C, Steinbrink C, Martin C, Wolf RC: Neural correlates of working memory performance in adolescents and young adults with dyslexia. Neuropsychologia 2008; 46:640–648.

67. Verius M, Frank F, Gizewski E, Broessner G (2019) Magnetic resonance spectroscopy thermometry at 3 tesla: importance of calibration measurements. Ther. Hypothermia Temp. Manag. 9, 146–155. doi: 10.1089/ther.2018.0027

68. Xu H, Robertson RM (1996) Neural parameters contributing to temperature compensation in the flight CPG of the Locust, Locusta migratoria. Brain Res. 734, 213–222. DOI: 10.1016/0006-8993(96)00635-X

69. Yablonskiy DA, Ackerman JJH, Raichle MF (2000) Coupling between changes in human brain temperature and oxidative metabolism during prolonged visual stimulation. PNAS, 97, 7603–7608. doi: 10.1073/pnas.97.13.7603

70. Yeganeh AJ, Reichard G, McCoy AP, Bulbul T, Jazizadeh F (2018) Correlation of ambient air temperature and cognitive performance: A systematic review and meta-analysis. Building and Environment 143, 701–716. doi: 10.1016/j.buildenv.2018.07.002

71. Zhang Y, Taub E, Mueller C., Younger J, Uswatte G, DeRamus TP, Knight DC (2020) Reproducibility of whole-brain temperature mapping and metabolite quantification using proton magnetic resonance spectroscopy. NMR in biomedicine, 33(7), e4313. 10.1002/nbm.4313

72. Zecevic D, Levitan H (1980) Temperature acclimation: effects on membrane physiology of an identified snail neuron. Am. J. of Physiol. - Cell Physiol. 239, C47–C57. doi: 10.1152/ajpcell.1980.239.3.C47

